# Large-Scale Assessment of *NF1* Single Amino Acid Variants as HLA Class I Neoantigens

**DOI:** 10.64898/2026.05.10.724138

**Authors:** Sung Yun Jung, Aylar Babaei, Alexandros Tzatsos, Junfeng Ma, Yang Yu, Wai Chin Chong, Huizhen Zhang, Richard T. Graham, Conrad Russell Cruz, Javad Nazarian, Brian R. Rood, Jianhua Yang, Chunchao Zhang

**Affiliations:** Department of Molecular and Cellular Biology, Baylor College of Medicine, Houston, TX, USA; Center for Cancer and Immunology Research, Children’s National Research Institute, Children’s National Hospital, Washington, DC, USA; Department of Anatomy and Cell Biology, George Washington University Cancer Center, Washington, DC, USA; Department of Oncology, Lombardi Comprehensive Cancer Center, Georgetown University Medical Center, Georgetown University, Washington, DC, USA; Division of Oncology, Cancer and Blood Diseases Institute, Cincinnati Children’s Hospital Medical Center, Cincinnati, OH, USA; Department of Pediatrics, The George Washington University School of Medicine and Health Sciences, Washington, DC, USA; DIPG/DMG Center Zurich, Children’s Research Center, University Children’s Hospital Zurich, University of Zurich, Zurich, Switzerland; Department of Otorhinolaryngology - Head and Neck Surgery, The University of Maryland School of Medicine, Baltimore, MD, USA

**Author notes:** Correspondence to: C Zhang, Department of Otorhinolaryngology - Head and Neck Surgery, The University of Maryland School of Medicine, Baltimore, MD, USA. **Conflict of interest statement:** These authors declare no conflict of interests.

**Keywords:** Neurofibromatosis Type 1, Neoantigen, HLA binding prediction

## Abstract

Neoantigens are cancer-specific antigens arising from genomic alterations. Single Amino Acid Variants (SAAVs) represent a primary class of these neoantigens. To evaluate the therapeutic potential of Neurofibromin 1 (*NF1*)-derived SAAVs - given that *NF1* is frequently mutated in malignant brain tumors - we prioritized the 40 *NF1* SAAVs determined to be HLA-A*02:01 binders using computational prediction coupled with experimental validation. To validate these predicted neoepitopes, we employed a two-tiered experimental approach in HLA-A*02:01 homozygous U87-MG cells. We first synthesized minigene constructs encoding the predicted neoepitopes, introduced them via lentiviral transfection and confirmed their expression by mass spectrometry (MS). Subsequently, we performed endogenous validation using pan-HLA immunoprecipitation mass spectrometry (IP-MS), confirming 4 (10 neoepitopes) of the 40 candidate SAAVs. We observed a discrepancy between *in silico* predictions and the observed sequences. Our endogenous peptidomics further revealed conserved peptide motifs and demonstrated that peptide selection for HLA presentation is transient. While our study substantiates the therapeutic feasibility of T-cell immunotherapies targeting *NF1* mutations, these results underscore a limitation in current computational prediction. Our study highlights the necessity of experimental validation to refine neoantigen prioritization strategies.

## 1. Introduction

Neoantigens are cancer-specific peptides that arise from cancer-causing gene alterations[1]. As neoantigens are absent in normal tissues, they could be recognized as non-self by the host T-cell-mediated adaptive immune system[2]. Neoepitopes, the antigenic determinants of neoantigens, are presented on Major Histocompatibility Complex (MHC) molecules as a peptide-MHC (pMHC) complex. The pMHC complex is then recognized by T cells carrying cognate T cell receptors (TCRs) to trigger a T cell-mediated immune response that attacks cancer cells. Thus, the peptide-MHC-TCR (pMHC-TCR) interaction plays a central role in mounting a powerful host immune response. To tackle many non-self-threats, the multimeric pMHC, TCR, and pMHC-TCR complexes are equipped with many rearrangeable components, ensuring broad diversity and wide antigen coverage. Since most cancer-specific genetic alterations are unique to an individual patient and generate exclusive neoantigens, the response of the neoepitope-MHC-TCR complex is highly specific, making them an ideal platform for developing personalized cancer immunotherapies.

The Neurofibromin 1 (*NF1*) gene is a critical tumor suppressor located on chromosome 17q11.2 and has a length that spans 60 exons[3, 4]. It encodes neurofibromin, a GTPase-regulating protein that regulates the RAS-signaling pathway, which is essential for controlling cell growth and proliferation[5]. *NF1* is frequently mutated in brain cancers[6]. Pathogenic mutations in the *NF1* gene can lead to an autosomal dominant genetic disorder known as Neurofibromatosis Type 1. This syndrome predisposes individuals to various tumors and has a global prevalence estimated to affect 1 in 2,000 to 1 in 5,000 people[7–9]. Within the central nervous system (CNS), *NF1* mutation is associated with infiltrating glioma, optic pathway glioma, and astrocytoma with piloid features[10–12].

To evaluate the therapeutic potential of *NF1*-derived SAAVs, we selected the 40 *NF1* SAAVs predicted as HLA-A*02:01 binders by using two computational algorithms. To experimentally validate these predicted neoepitopes, we designed eight constructs, each containing five minigenes cover five SAAVs predicted as neoepitopes. We synthesized the constructs and introduced them into U87-MG cells via lentiviral transfection, as these cells highly express homozygous HLA-A*02:01 - one of the most common HLA Class I alleles across different races[13]. We confirmed the expression of seven constructs by mass spectrometry (MS). To evaluate the expression and peptide length in the proteasome stage prior to antigen presentation, we performed MS analysis of endogenous peptides in the cells excluding the intact proteins. Subsequently, we performed pan-HLA immunoprecipitation mass spectrometry (IP-MS) and validated 4 of the 40 SAAVs or 10 neoepitopes as HLA-A*02:01 binders. A discrepancy between *in silico* predictions and the detected sequences was observed. Our study underscores a limitation in current computational prediction and highlights the necessity of experimental validation to refine the therapeutical strategies targeting mutation derived neoantigens.

## 2. Materials and Methods

### 2.1 Cell culture

The human glioblastoma cell line U87-MG was obtained from the American Type Culture Collection (ATCC) and used for neoantigen validation. The HEK293T cell line was obtained from ATCC and used for lentivirus packaging. U87-MG cells were grown in Eagle’s Minimum Essential Medium (EMEM) supplemented with 10% Fetal Bovine Serum (FBS) and 1% Penicillin-Streptomycin. HEK293T cells were grown in DMEM containing 10% heat-inactivated fetal bovine serum (Invitrogen), 100 units/mL penicillin, and 100 µg/mL streptomycin (Invitrogen).

### 2.2 Minigene constructs and lentiviral transduction

We designed eight minigene constructs for experimental validation of predicted *NF1* neoepitopes. Each construct has five minigenes and each minigene encodes a predicted neoepitope along with 12 flanking amino acids on either side. Sequences are separated by a glycine-serine (GS) linker. The full construct also includes a signal peptide, Kozak sequence, and MHC-I trafficking signal. Synthetic DNA constructs, approximately 1 kb in length, were produced by Twist Bioscience (South San Francisco, CA).

The synthetic DNA constructs were cloned into a lentiviral expression vector with blasticidin resistance for selection. HEK293T cells were used to package the lentiviruses, which were then used to infect U87-MG cells - a glioblastoma line with homozygous HLA-A*02:01 and high HLA expression. For lentivirus production, 2.5 × 10⁶ HEK293T cells were seeded in a 10-cm dish 24 h prior to transfection. Cells were transfected using GeneJuice® Transfection Reagent (Cat. #70967, MilliporeSigma) with 6 µg of lentiviral vector, 2 µg of psPAX2 packaging plasmid (Cat. #12260, Addgene), and 2 µg of pMD2.G envelope plasmid (Cat. #12259, Addgene). Lentivirus-containing supernatant was collected 48 h post-transfection. U87-MG cells were transduced in the presence of polybrene (4 µg/mL) and subsequently selected with blasticidin (8 µg/mL; MilliporeSigma) for 5 days.

### 2.3 Cellular proteome

Construct expression in U87-MG cells was confirmed by MS. Briefly, around 1×10^6^ U87-MG cells transduced with the constructs were collected by centrifugation, washed with cold PBS, and subsequently processed using the S-Trap™ system (Protifi, NY, USA). The cell pellet was resuspended and lysed in 5% SDS and 50 mM TEAB (pH 8.5) via sonication. Protein concentration measured by BCA kit and 20 ug of proteins were reduced with 120 mM TCEP and alkylated using 500 mM MMTS in isopropanol. Following acidification with phosphoric acid, proteins were precipitated by the addition of 100 mM TEAB in 90% methanol. The precipitated proteins were loaded onto the S-Trap device and digested overnight with 1 ug of Trypsin/Lys-C (A41007, Thermo Scientific). Resulting peptides were recovered, vacuum-dried, and quantified using a Colorimetric Peptide Assay kit (23275, Thermo Scientific). For each run, 500 ng of peptides were analyzed on a Vanquish HPLC system coupled to an Orbitrap Astral mass spectrometer (Thermo Scientific) equipped with a 15 cm PepMap™ RSLC C18 analytical column and an electrospray ionization source. For data-dependent acquisition (DDA), an Orbitrap lumos (Thermo Scientific) mass spectrometer was utilized. A top 3 second method was used to select precursor ions from a full MS scan (400–1,400 m/z, 120,000 resolution) followed by HCD fragmentation. Alternatively, a timsTOF Ultra (Bruker) mass spectrometer was utilized under DDA mode. For data-independent Acquisition (DIA), an Orbitrap Astral mass spectrometer (Thermo Scientific) was used. MS1 scans were acquired in the Orbitrap at resolution of 240,000 (m/z 380–980) with EASY-IC™ internal calibration. DIA MS/MS spectra were collected across the same precursor range using 2 m/z isolation windows, HCD fragmentation (25% NCE), and detection in the Astral mass analyzer (m/z 150–2,000).

Obtained spectra were searched against the target-decoy human proteome database plus eight custom construct sequences by using fragpipe v24.0. The human proteome database (2023-03-01-decoys-reviewed-contam-UP000005640.fas within fragpipe) contains UniProt reviewed sequences, reversed sequences, contaminants, and the sequences of eight synthetic constructs. An LFQ-MBR.workflow in fragpipe was adopted for spectra acquired by DDA. A DIA_SpecLib_Quant.workflow in fragpipe was employed for spectra acquired by DIA. Cutoffs were 1% peptide FDR and 1% protein FDR. The whole proteome MS data were collected in triplicates.

### 2.4 Cellular Peptidome

To determine the expression and the peptide length in the proteasome stage prior to antigen presentation, we performed MS analysis of non-specific peptides in the cells excluding the intact proteins. Briefly, ∼1×10^6^ U87-MG cells transduced with the constructs were collected by centrifugation, washed with cold PBS, lysed in 8 M urea and filtered through 30K membrane filter (Sartorius, Vivaspin® 500) at 10,000 g at 4 °C. The filtrate was vacuum dried and rehydrate using 0.5% formic acid. Per MS run, 500 ng of peptide were measured and run in DIA or DDA mode by Orbitrap Astral mass spectrometer (Thermo Scientific) described previously.

Obtained spectra were searched against the target-decoy database described previously. For DDA data, a nonspecific-HLA.workflow in fragpipe was adopted. For DIA data, a nonspecific-HLA-DIA-Astral.workflow in fragpipe was used. These workflows search peptides with length varies from 8 to 25 amino acids (AA). Cutoffs were 0.1% peptide FDR. The peptidomes data were collected in duplicates.

### 2.5 Immunoprecipitation mass spectrometry (IP-MS)

We performed IP-MS to detect HLA binders by using anti-human MHC Class I (HLA-A, HLA-B, HLA-C) antibody (BioXcell, Catalog Number: BE0079, anti-human MHC Class I monoclonal antibody). Briefly, ∼0.2 mL of cell pellets (∼1.0 × 10^8^ cells) was lysed by sonication in a lysis buffer containing 0.1% NP-40 and then cleared by ultracentrifugation (100kg, 4C, 20 min). The supernatant was incubated with preconditioned anti-human MHC Class I antibody-conjugated NHS-agarose beads. The peptide-HLA complex was eluted from the beads using 10% acetic acid, and the peptides will be further purified with the C18 Sep-PAK SPE column. Purified immune peptides were subjected to a nanoLC-1200 (Thermo Scientific) coupled to Orbitrap Astral mass spectrometer (Thermo Scientific) with an ESI source. A C18 trap column (2 cm, 100 μm i.d.) and a separation column (5 cm, 150 μm i.d.) were used for peptide enrichment and separation. Spectra were acquired by either DDA or DIA for unbiased peptide detection. Field Asymmetric Ion Mobility Spectrometry (FAIMS) was applied specifically for DDA mode.

Obtained spectra were searched against the target-decoy database as described previously. For DDA with FAIMS, a nonspecific-HLA.workflow in fragpipe was adopted. For DIA without FAIMS, a nonspecific-HLA-DIA-Astral.workflow in fragpipe was used. Cutoffs were 0.1% peptide FDR. Only peptides with length varies from 7 to 15 amino acids were considered as HLA-A*02:01 binders. Immunopeptidomes data were collected in duplicates.

### 2.6 Computational prediction of *NF1* SAAVs as neoantigens

To assess the feasibility of computationally predicting HLA-presented neoantigens, we analyzed *NF1* mutations in The cBioPortal for Cancer Genomics (cBioPortal) and Cancer Cell Line Encyclopedia (CCLE) data portals covering 9,603 brain tumors and 1,100 cancer cell lines respectively. A total of 156 unique *NF1* single nucleotide variants (SNVs) including missense and nonsense mutations (145 from cBioPortal glioma patient samples and 11 from CCLE glioma cell lines) were found in 226 *NF1* glioma samples.

Two HLA binding prediction algorithms, NetMHCpan 4.1 EL and consensus, on The Immune Epitope Database (IEDB) (https://www.iedb.org/home_v3.php) were adopted for binding prediction on HLA-A*02:01. Peptide length varies from 8 to 14 are considered and the best ranked epitopes covering SAAVs are selected. Among 156 unique *NF1* SAAVs, we selected 40 ranked in the top 10% for HLA-A*02:01 binding by NetMHCpan 4.1 EL[14] or the top ∼30% by consensus[15].

### 2.7 Bioinformatics analyses

U87-MG cell line was selected for experimental validation due to its high expression of HLA-A and homozygous HLA-A*02:01. RNA sequencing of U87-MG cells (GEO accession: GSE217347) confirmed HLA-A (gene ID: 3105) high expression with >200 FPKM (Fragments Per Kilobase per Million) measured in the wide-type cells, ranking at top 1% of all measured genes. The bioinformatics software, seq2HLA[16], was used to determine U87-MG HLA alleles, which confirmed homozygous HLA-A*02:01.

For motif analysis, Seq2Logo-2.0 (https://services.healthtech.dtu.dk/services/Seq2Logo-2.0/) was used for amino acid enrichment and depletion[17]. Parameters are: Logo type: Shannon, clustering method: Hobohm1, threshold for clustering (Hobohm1): 0.63, pseudo counts: 200, content units: Bits.

## 3. Results

### 3.1 Experimental design and general statistics

As a proof-of-concept that SAAVs can be computationally predicted and utilized as neoantigens, we investigated all *NF1* mutations in the cBioPortal and CCLE data portals and selected the 40 *NF1* SAAVs predicted to be HLA-A*02:01 binders by two prediction algorithms – NetMHCpan 4.1 EL and consensus (**sTable 1**). We generated eight stable cell lines that were transduced with an *NF1* construct expressing *NF1* SAAVs by lentivirus infection (**Fig. 1A**).

**Figure 1.**
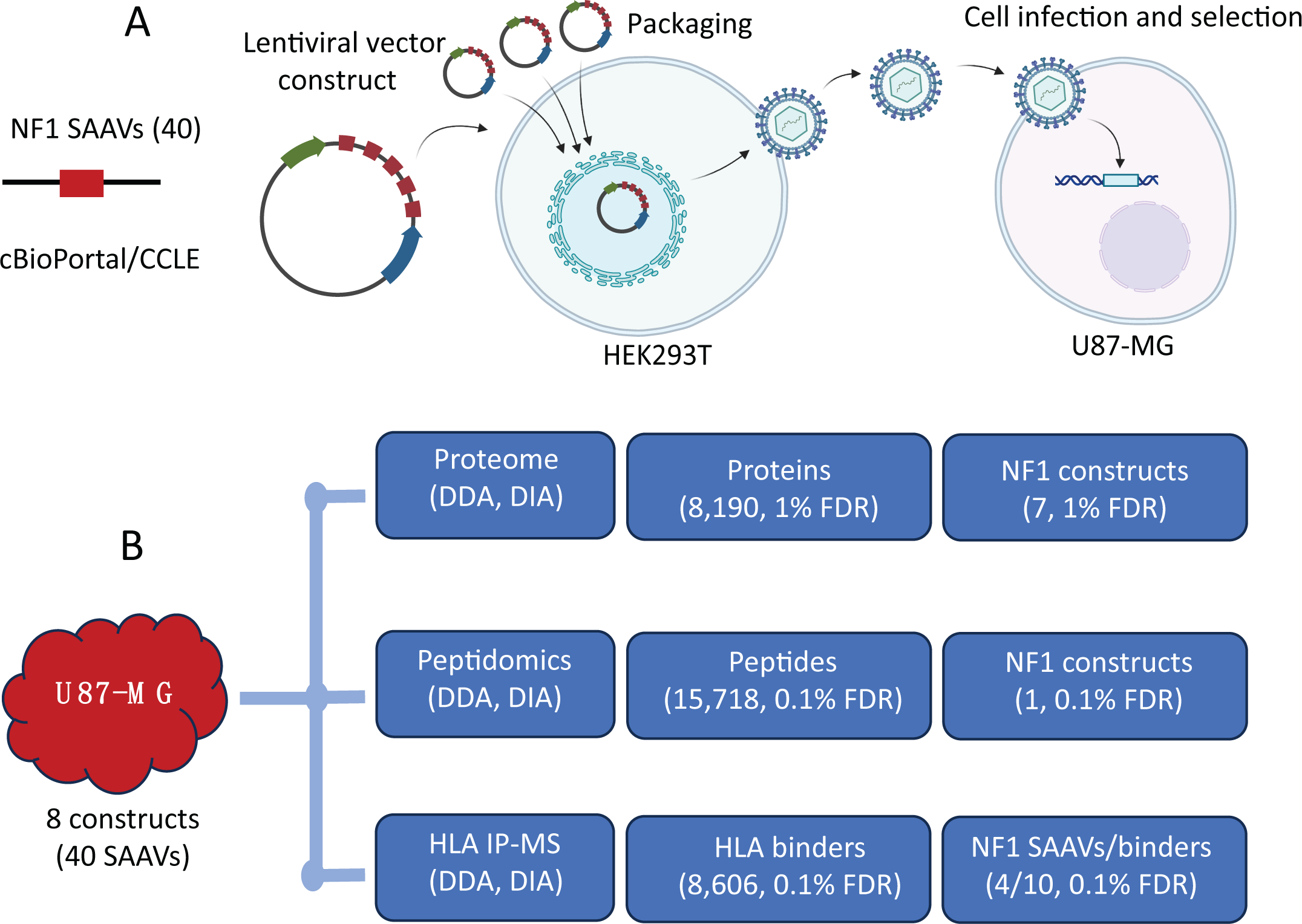
Experimental design and general statistics. (A) Schematic of minigene constructs and lentiviral transduction workflow. Forty *NF1* SAAVs were selected and cloned into a lentiviral vector, packaged in HEK293T cells, and stably transduced into U87-MG cells. (B) Mass spectrometry identification of *NF1* SAAVs, non-specific cytosolic peptides, and HLA-bound peptides.

LC-MS/MS was used to confirm the expression of *NF1* constructs by profiling the cellular proteome of 8 *NF1*-transduced U87-MG cell lines. Expression of 7 out of 8 *NF1* constructs was confirmed by identification of the construct’s unique peptides using different instruments and acquisition strategies (**Fig. 1B, sTable 2**). In the proteome analysis, a total of 8,190 proteins were identified with a 1% FDR at peptide and protein levels. On average, we observed ∼26% sequence coverage with at least one unique peptide detected for each construct (**Table 1**). As no pre-fractionation was used prior to LC-MS/MS, the total protein depth may have been reduced; we identified approximately 3,000 to 8,000 proteins per run. The identification of constructs and sequence coverage could likely be improved if a pre-fractionation strategy was adopted. The endogenous peptidome was explored to reveal the cytosolic peptides prior to antigen presentation. Among 15,718 peptides, we only detected one peptide from *NF1* constructs (**Fig. 1B, sTable 3**), suggesting that peptides in the pre-presentation stage are low-abundant and transient. To validate the computational prediction of HLA binders, pan-HLA-IP-MS identified a total of 8,606 HLA binders, 10 of which were validated as *NF1* neoepitopes representing 4 *NF1* SAAVs (**Fig. 1B, sTable 1 and 4**).

**Table 1.** Mass spectrometry-based validation of *NF1* minigene expression across all constructs.

| **Construct ID** | **Peptide Length** | **Sequence Coverage** | **Total Peptides** | **Unique Peptides** | **Instrument** | **Acquisition** |
| --- | --- | --- | --- | --- | --- | --- |
| NF1_C1 | 314 | 33.44 | 7 | 5 | Astral | DIA |
| NF1_C1 | 314 | 41.4 | 8 | 4 | TimsTOF | DDA |
| NF1_C2 | 322 | 8.39 | 2 | 1 | Astral | DIA |
| NF1_C2 | 322 | 15.22 | 3 | 1 | TimsTOF | DDA |
| NF1_C3 | 321 | 42.06 | 8 | 6 | Astral | DIA |
| NF1_C3 | 321 | 33.96 | 6 | 4 | Lumos | DDA |
| NF1_C3 | 321 | 50.47 | 10 | 7 | TimsTOF | DDA |
| NF1_C5 | 326 | 13.19 | 2 | 1 | TimsTOF | DDA |
| NF1_C6 | 317 | 11.99 | 1 | 1 | Lumos | DDA |
| NF1_C7 | 320 | 30.94 | 7 | 4 | Astral | DIA |
| NF1_C7 | 320 | 6.88 | 1 | 1 | Lumos | DDA |
| NF1_C7 | 320 | 37.81 | 7 | 4 | TimsTOF | DDA |
| NF1_C8 | 321 | 9.66 | 2 | 1 | Astral | DIA |

### 3.2 *NF1* SAAVs are ideal neoantigen candidates for cancer immunotherapy

We systematically investigated *NF1* mutations in the cBioPortal and CCLE data portals, which together cover 9,603 brain tumors and 1,100 cancer cell lines representing the most comprehensive catalogue of *NF1* mutations in gliomas. A total of 156 unique *NF1* SNVs including missense and nonsense mutations (145 from cBioPortal glioma patient samples and 11 from CCLE glioma cell lines) were found in 226 *NF1* gliomas. The resulting protein sequence alterations were randomly distributed across all 2,839 amino acids, with no preference for any known functional domains (**Fig. 2A**).

**Figure 2.**
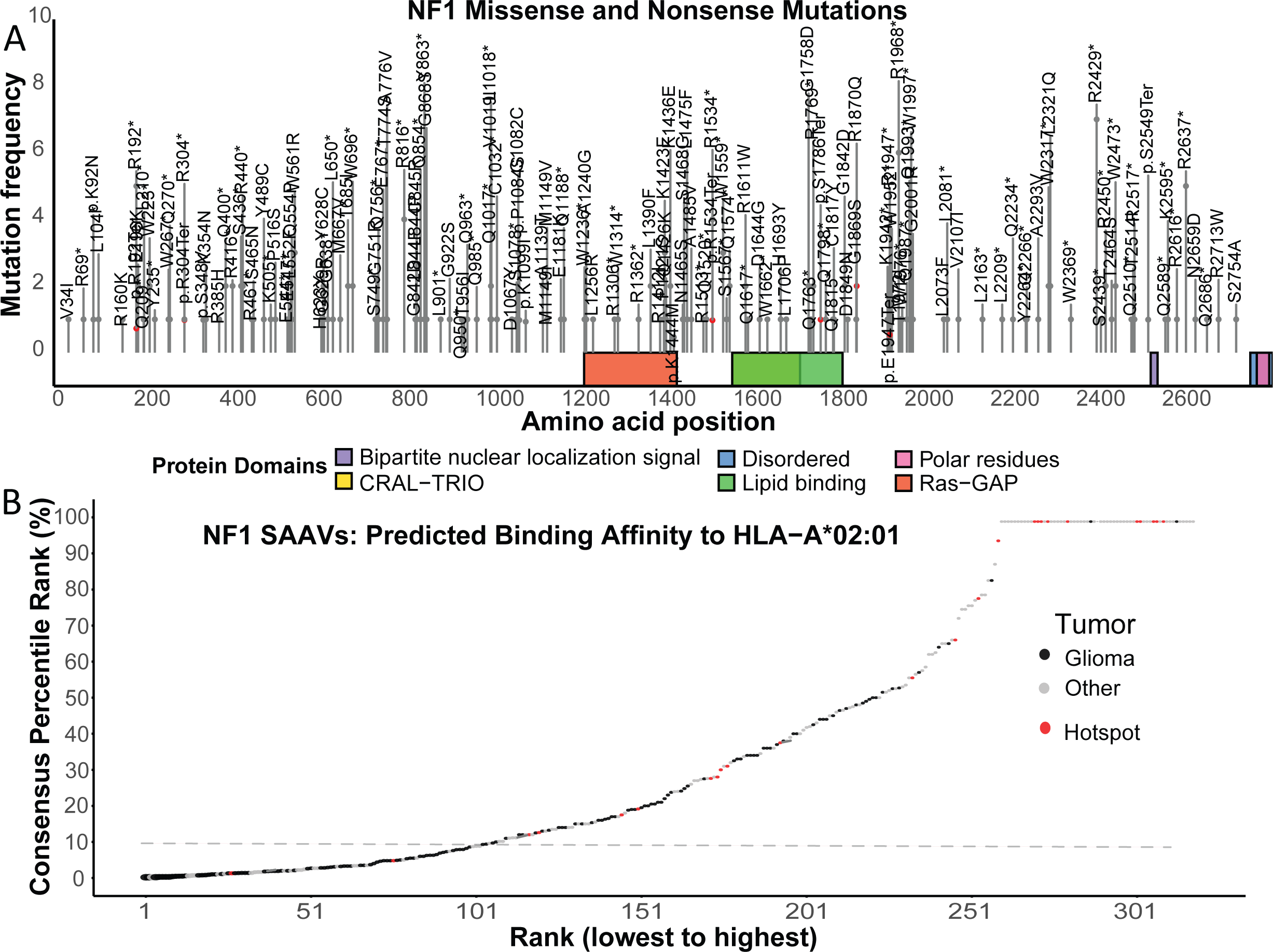
Computational evaluation of *NF1* SAAVs as potential neoantigens. (A) Mapping the distribution of *NF1* missense and nonsense mutations across the protein functional domains. Functional domains are indicated by colored blocks. (B) *NF1* SAAVs predicted as HLA-A*02:01 binders using the IEDB consensus algorithm. The horizontal dashed line represents the 10% percentile ranking threshold. Black dots are mutations identified in gliomas; grey dots are mutations from other CNS tumors; Red dots are hotspot mutations curated from the cBioPortal and CCLE databases.

We further extended *NF1* mutations to CNS tumors in the two data portals. We evaluated the MHC-I binding affinity of 318 unique SNVs on HLA-A*02:01 by the prediction algorithm consensus on The Immune Epitope Database (IEDB). Consensus percentile rank was calculated by using four different scoring matrix methods[15]. On HLA-A*02:01, 107 *NF1* mutations (∼34%) were within the top 10% consensus rank (**Fig. 2B**) demonstrating the potential for developing TCR-T cell therapies targeting *NF1* mutations as cancer-specific antigens.

### 3.3 Peptides in the cytosol prior to antigen presentation

HLA class I molecules present a broad spectrum of peptides to cytotoxic T lymphocytes, allowing the immune system to recognize intracellular pathogens. Antigenic peptide precursors are produced by the proteasome and further trimmed by cytosolic aminopeptidases. They are translocated into the endoplasmic reticulum (ER) lumen by the peptide transporter TAP (Transporter associated with Antigen Processing) and are further trimmed in the ER lumen by ER aminopeptidases before selection by a defined MHC class I allele[18]. Only a limited number of peptides are selected and presented by a given HLA allele. How a defined MHC I allele selects the correct peptides for presentation out of a large and diverse peptide pool is largely unknown.

Our peptidomic experiments identified a total of 15,718 non-specific peptides at 0.1% peptide FDR (**Fig. 1B, sTable 3**). The length of these peptides varies from 8 to 25 AA and majority fall in a range of 9 to 16 AA (**Fig. 3A**). Motif analyses of the 9- to 16-mer peptides revealed conserved residues in the first and last position (**Fig. 3B-I**). Enrichment of Alanine (A) and Serine (S) at the first position, and Arginine (R), Leucine (L), and Lysine (K) at the last position are observed regardless of the peptide length. The N-terminal alanine enrichment suggests a preferential trimming mechanism mediated by alanine aminopeptidases during the maturation of cytosolic peptide precursors[19], while the enrichment of R, L, and K at the peptide C-termini suggests processing by cytosolic trypsin-like and chymotrypsin-like proteases[20]. These enzymes preferentially cleave after basic (K, R) and hydrophobic (F, Y, W, L) residues, respectively.

**Figure 3.**
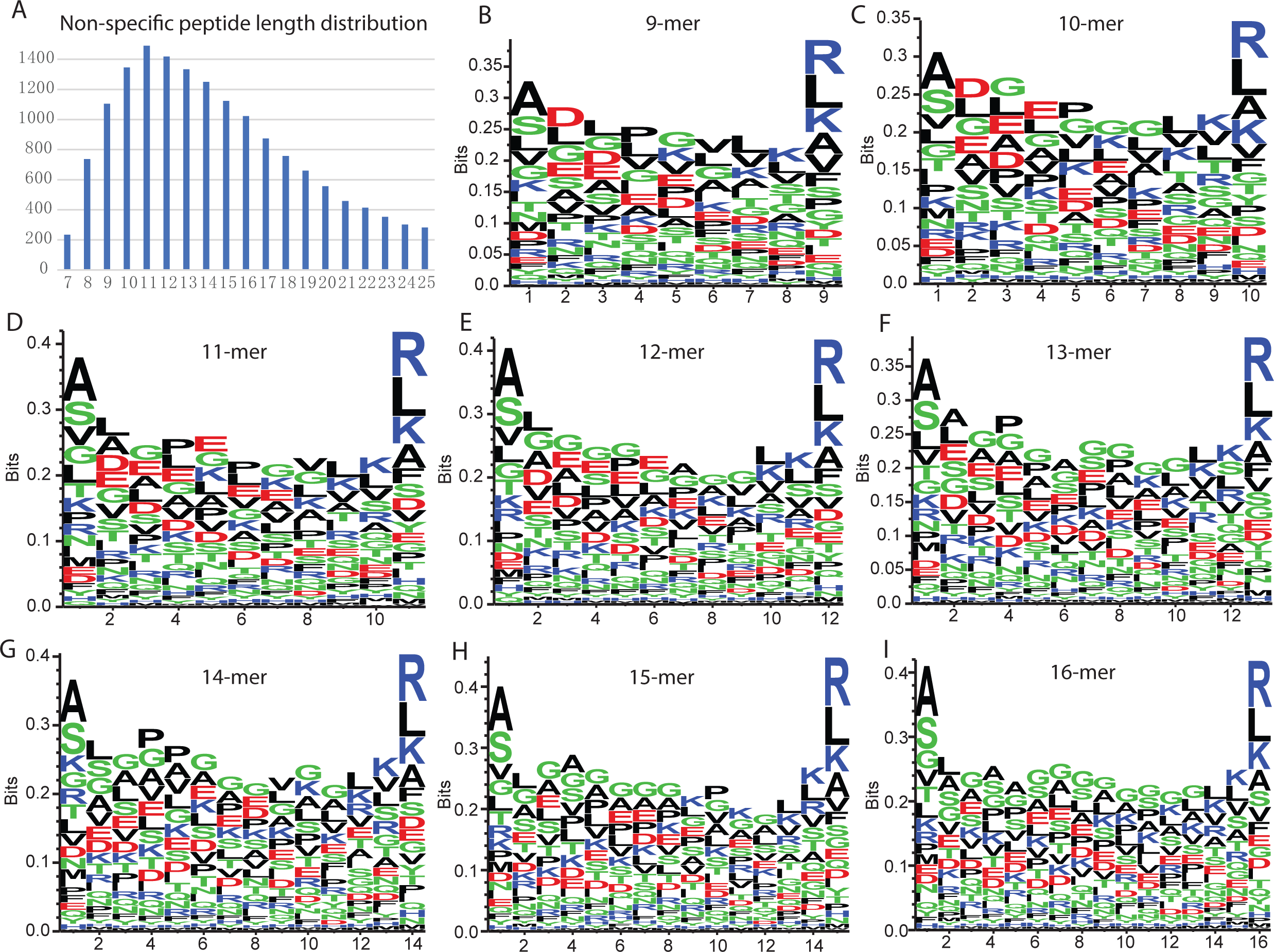
Characterization of non-specific cytosolic peptides. (A) Length distribution of non-specific peptides identified by MS. (B–I) Sequence motif analysis of non-specific peptides categorized by length, ranging from 9- to 16-mers.

Compared to the construct-specific peptides identified in cellular proteome (**Table 1**), we only detected six peptides mapped to *NF1* gene and one peptide uniquely mapped to the NF1_C7 construct (**sTable 3**), suggesting that the degradation of these constructs in the proteasome is rapid and transient.

### 3.4 Antigen presentation by HLA class I molecules

We conducted pan-HLA-IP-MS to identify HLA class I binders in U87 cells. At 0.1% peptide FDR, a total of 8,606 HLA class I epitopes were identified including 10 neoepitopes covering 4 SAAVs (**Fig. 1B, sTable 1 and 4**). Most of the HLA binders fell in a range of 8 to 11 AA. The 9-mer peptides are predominant as expected (**Fig. 4A**). Motif analysis of the 9-mer peptides revealed conserved residues - Leucine (L) and Glutamic acid (E) - at the second position, and residues including Leucine (L) and Valine (V) at the ninth position, which is expected and consistent with the known binding motif of the HLA-A*02:01 allele (**Fig. 4B**).

**Figure 4.**
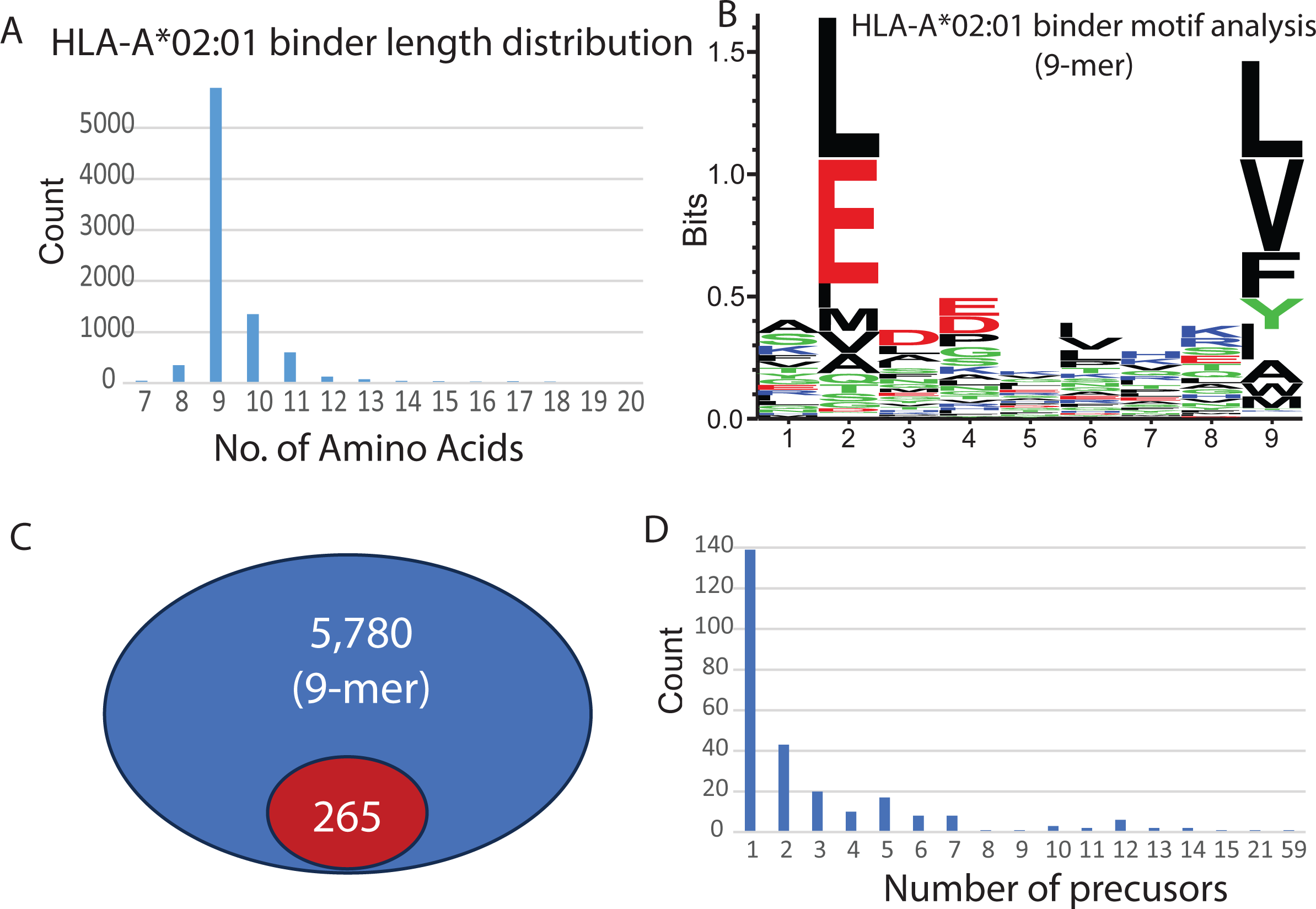
Profiling and precursor mapping of HLA-bound peptides identified via IP-MS. (A) Length distribution of HLA-A*02:01 binders identified by immunopeptidomics. (B) Sequence motif analysis of identified 9-mer HLA-A*02:01 binders. (C) Number of 9-mer HLA-A*02:01 binders identified with precursors in the non-specific cytosolic peptide pool. The number in the blue oval represents the total count of identified 9-mer HLA-A*02:01 binders, while the number in the red oval indicates the subset of these binders for which corresponding precursors were detected in the non-specific cytosolic peptide pool. (D) Distribution of the number of cytosolic precursors identified for each HLA-A*02:01 binder.

To characterize the precursors before they are presented by HLA, we chose 5,780 9-mer peptides and searched for their precursors in the endogenous peptidomics prior to HLA presentation (**Fig. 4C**). We searched peptides whose sequences are completely contained within the endogenous peptide set (minimum length of 9). Among the 5,780 binders, only 265 (4.6%) were found among the endogenous cytosolic peptides. Most of them (139) were found in one peptide precursor, however 126 binders were found in multiple precursors (**Fig. 4C, sTable 5**). 18 binders were detected with more than 10 precursors. The sequence, SLGSALRPS, is part of the Vimentin protein. It had 59 identified precursors ranging from 9 to 25 amino acids (**sTable 5**), suggesting that for a given HLA allele, the presented peptide is precisely selected from a large peptide pool. Most of the binders were not found in a precursor suggesting rapid protein degradation by cytosolic aminopeptidases.

### 3.5 Computational prediction and experimental validation of *NF1*-associated neoantigens in brain tumors

To assess the feasibility of computationally predicting HLA-presented neoantigens, we analyzed *NF1* mutations, one of the most frequently mutated genes in brain tumors, from 226 gliomas in cBioPortal and CCLE. Among 156 unique *NF1* SNVs, we selected 40 *NF1* SAAVs with a best score ranked in the top 10% for HLA-A*02:01 binding by NetMHCpan 4.1 EL[14] or the top ∼30% by consensus[15] (**sTable 1**). We designed eight minigene constructs as previously with some modifications[21], each containing five SAAVs separated by GS linkers (**Fig. 5A, sFig. 1**). Each minigene encoded a predicted neoepitope along with 12 flanking amino acids on either side (**Fig. 5A**). The constructs also included a signal peptide for secretion, a Kozak sequence, and an MHC-I trafficking signal (**Fig. 5A, sFig. 1**). The constructs were cloned into a lentiviral expression vector with blasticidin resistance for selection. HEK293T cells were used to package the lentiviruses, which were then used to infect U87-MG cells - a glioblastoma line with homozygous HLA-A*02:01 and high HLA expression. Expression of the seven constructs was confirmed by MS (**Table 1**).

**Figure 5.**
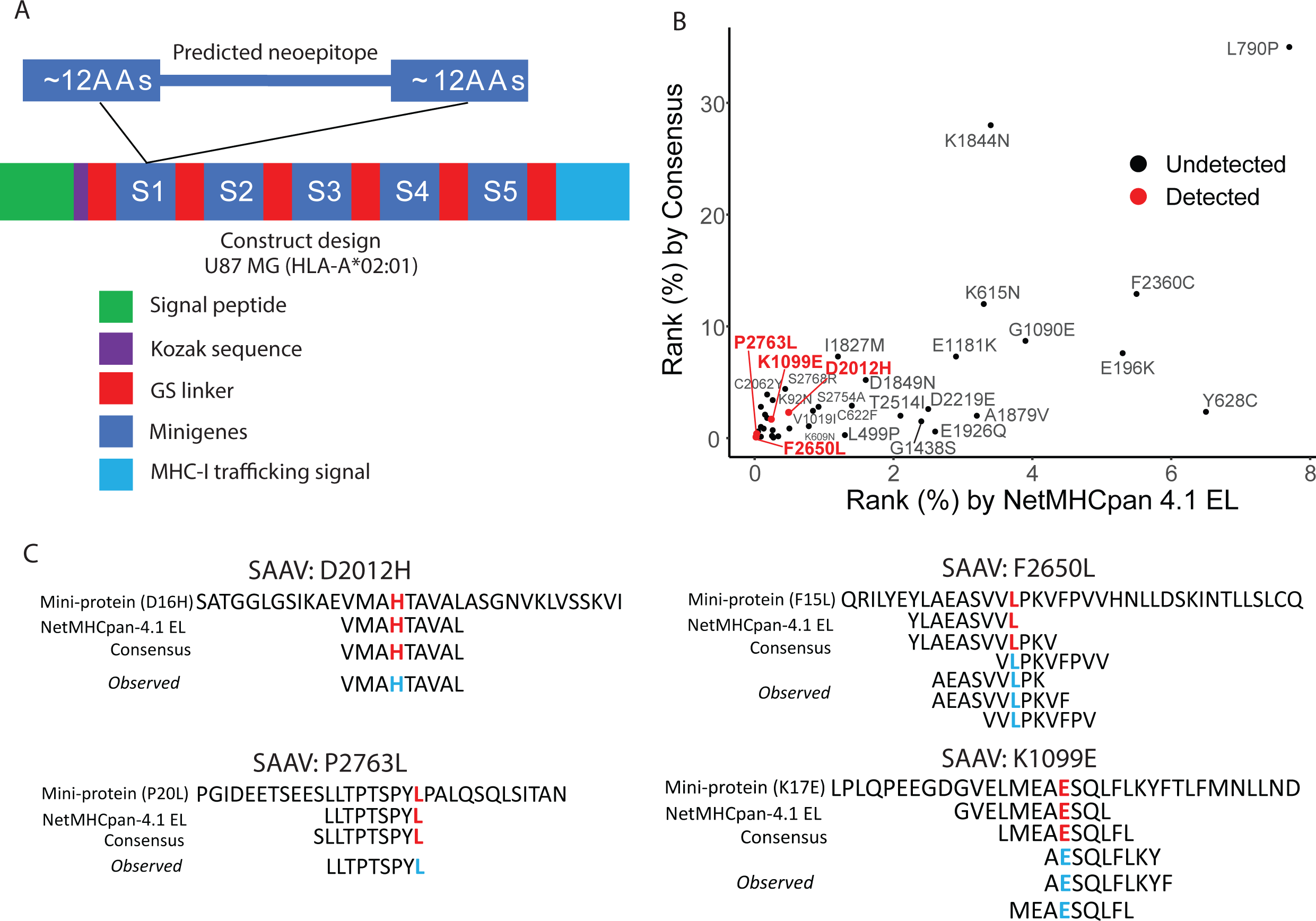
Experimental validation of *NF1*-associated neoantigens. (A) Schematic details of eight minigene constructs. SAAVs are separated by flexible glycine-serine (GS) linkers. Each minigene encodes a predicted neoepitope flanked by 12 amino acids on both the N- and C-termini to ensure authentic proteolytic processing. The constructs incorporate a Kozak sequence for translation initiation, a signal peptide for secretion, and an MHC-I trafficking signal (MITD) to facilitate efficient antigen presentation. (B) Percentile rank distribution of 40 SAAVs by two prediction algorithms, with the successful detection of four specific SAAVs by IP-MS highlighted in red. (C) Alignment and comparison of observed neoepitope sequences against predicted sequences from each algorithm. Residues in red or cyan font indicate the position of the SAAV within the respective sequences, categorized by minigene-encoded sequences, predicted sequences, and MS-observed sequences.

To validate predicted HLA-A*02:01 neoepitopes, we performed pan-HLA-IP-MS and identified 10 neoepitopes across four distinct SAAVs (**Fig. 5B, sTable 1 and 4**). While several high-ranking SAAVs remained undetected, all MS-validated SAAVs were predicted to be within the top 1% by NetMHCpan 4.1 EL or the top 3% by the consensus method, highlighting both the potential and current limitations of computational neoantigen prediction (**Fig. 5B**). Notably, the predicted sequences for these four SAAVs did not always align with the observed peptides (**Fig. 5C**). For example, the D2012H variant (Aspartic acid to Histidine at position 2012), both algorithms correctly predicted the observed sequence. However, the P2763L variant (Proline to Leucine at position 2763), the observed sequence matched the NetMHCpan-4.1 EL prediction specifically, while the consensus method varied slightly. For the remaining SAAVs, we observed multiple peptide forms per variant that differed from the algorithmic predictions. For instance, five unique sequences were detected for F2650L (Phenylalanine to Leucine at position 2650) and three for K1099E (Lysine to Glutamate at position 1099), none of which were exactly captured by the two algorithms. These findings suggest that neoepitopes often exist as multiple isoforms and highlight the need for improved predictive accuracy.

## 4. Discussion

The identification of cancer-specific antigens is a cornerstone of personalized immunotherapy, yet the vast majority of structural variants remain ‘dark’ to current predictive algorithms. In this study, we provide a robust proof-of-concept for the systematic validation of SAAVs as actionable targets. By integrating lentiviral minigene expression with high-stringency pan-HLA-IP-MS, we have demonstrated that while computational tools may have a role for peptide prioritization, they do not capture the full complexity of antigen processing and presentation.

Expression of seven lentiviral constructs in U87-MG cells was validated through proteomic profiling. Using different MS and data acquisition strategies, we confirmed stable expression, thereby establishing the robustness of our synthetic design. For the cellular proteome analysis, we utilized a simplified sample preparation workflow without pre-fractionation. Although this resulted in a lower protein yield, the successful detection of our targets under these conditions suggests that the implementation of offline pre-fractionation could further improve the MS detectability.

A significant number of *NF1* SAAVs are predicted to be candidate neoepitopes, ranking within the top 10% for the consensus method. However, the distribution of *NF1* SAAVs is largely random and sporadic across all known *NF1* protein domains and it is a very large gene. Since hotspot SAAVs are rare, it may be advantageous to pursue a personalized approach targeting “private” neoantigens.

Our endogenous peptidomics revealed that the majority of these cytosolic peptides range from 9 to 16 AA in length. For the first time, our motif analysis identified conserved residues at the first and last positions that are length-independent. Our results also suggest that cytosolic peptides are transient, low-abundance, and rapidly degraded by cytosolic proteases prior to HLA antigen presentation. This may explain why certain *NF1* SAAVs, predicted as HLA-A*02:01 binders within the top 1% by NetMHCpan 4.1 EL, remain undetected by IP-MS as protein degradation in the cytosol is highly heterogeneous. Consequently, this suggests that precursor stability and turnover rates should be considered into future computational prediction models.

We developed a robust immunopeptidomics approach, identifying a total of 8,606 HLA binders in U87-MG cells at a 0.1% peptide FDR - a significantly more stringent cutoff than the 1% FDR typically used in conventional proteomics. Approximately 67% (5,780) of these binders were 9-mer sequences. The presence of expected conserved residues at the second and ninth positions further demonstrates the reliability and robustness of our pan-HLA-IP-MS pipeline. Notably, only 265 (4.6%) of these 9-mer HLA binders had detectable precursors in the cytosol, highlighting the challenge of capturing transient neoepitope precursors. While most HLA binders originated from a single precursor, some were identified with more than ten, suggesting that the selection of peptides for HLA presentation from the total cytosolic pool is highly selective and precise.

Current computational algorithms have limitations in predicting HLA binders. Many peptides predicted to be high-affinity binders by software were not actually detected by MS, highlighting the “prediction gap” in current bioinformatics. Among the 40 selected *NF1* SAAVs, only four were detected by pan-HLA-IP-MS. Furthermore, the predicted sequences did not always align with the observed ones, suggesting that top-ranked predictions may not represent the optimal sequences selected by a given allele for surface presentation. Interestingly, two SAAVs were associated with multiple detected neoepitopes, indicating that the same variant can be presented in multiple isoforms.

This gap was also observed in a previous study[22]. Among thousands of SAAVs determined by whole exome sequencing, only 11 neoepitopes were identified by IP-MS in five patients and none of the identified neoepitopes predicted as top-ranked binders by NetMHCPan using a threshold of <500 nM as predicted affinity[22]. Most prediction algorithms are trained on static snapshots of eluted ligands or *in vitro* binding affinity data. This may effectively catch the conserved sequence motifs; however, the dynamic, multi-step processing pathway that occurs within the cytosol is ignored. Our study underscores the need for improved predictive accuracy and the necessity of experimental validation when selecting neoepitopes for T-cell immunotherapies.

The current study has limitations. First, while MS-based approach remains the most powerful tool for high-through immunopeptidomics, the issue of the sensitivity is the major challenge for detecting the endogenous, low-abundance neoantigens derived from low-abundance transcripts. Second, our validation workflow was limited to the U87-MG cell line. Future studies evaluating the expression of *NF1* SAAVs in a broader range of host cells - particularly professional antigen-presenting cells (APCs) with matched HLA alleles - are essential to ensure the generalizability and robustness of our findings across different cellular environments. Third, there is an MHC-I signal peptide in our design but whether it impacts the propensity of the peptides to go to the surface is unclear. Like in the natural processing state, this signal peptide may be absent in the future design. Fourth, as our selection of 40 *NF1* SAAVs focused on HLA binding, the subsequent identification of neoantigen-specific T cells and the evaluation of T-cell cytotoxicity are essential next steps for the clinical development of T-cell-based immunotherapies. Finally, while the utilization of a homozygous HLA-A*02:01 model provided the experimental stringency required for this proof-of-concept, the extensive allelic polymorphism within the human population means a vast landscape of ‘dark’ variants associated with non-A2 supertypes remains uncharacterized.

## Acknowledgements

This research was partially Supported by Grant #IRG 22-973396 from the American Cancer Society and from the Katzen Foundation to CZ, and the NIH/NINDS grant 5R01NS118008 to JY.

## Author contribution statement

SJ, AT, and CZ designed the study. YY, JM, WC, and HZ carried out experiments. AB, SJ and CZ analyzed the data. SJ, AT, JY, and CZ interpreted the results of the experiments. SJ and CZ drafted the manuscript. RG, CC, JN, BR, JY edited the manuscript. All the authors approved the final version of the manuscript.

## Data availability statement

Mass spectrometry data and associated files were deposited to the MassIVE repository and are accessible under the identifier MSV000101764.

## Declaration of generative AI use

Gemini was used only for language editing.

