## Supplemental Legends for "Large-Scale Assessment of *NF1* Single Amino Acid Variants as HLA Class I Neoantigens"

**Supplementary Figure 1. Detailed amino acid sequences and its corresponding nucleotide sequences of eight constructs.** Yellow highlights: 5' or 3' linkers. Green text: Kozak sequences (GCCACC). Green highlights: MHC class I signal peptides. Cyan highlights: MHC class I trafficking signal (MITD). Red and pink text: flexible glycine-serine (GS) linkers. Underlined text: minigene segments encoding predicted neoepitopes.

**Supplementary Tables:**

**Table S1. Detailed summary of the 40 *NF1* single amino acid variants (SAAVs) selected for experimental validation.** Data include genomic coordinates, cDNA and amino acid substitutions, and extended flanking sequences. Computational metrics are percentile ranks across two prediction algorithms. The "Detected sequence" column denotes specific neoepitopes successfully identified and confirmed via HLA IP-MS.

**Table S2. A list of all proteins identified by LC-MS/MS.** Stringent filtering criteria were applied to ensure high-confidence identifications: a Protein False Discovery Rate (FDR) of <1%, a Peptide FDR of <1%, and a requirement of at least one unique peptide per protein. All entries were filtered against the decoy and contaminants databases.

**Table S3. Comprehensive profiling of the non-specific cytosolic peptidome via LC-MS/MS. Sheet 1:** A list of all peptides identified by different instruments and acquisition strategies. **Sheet 2:** Non-redundant peptides and length distribution. **Sheet 3:** *NF1*-Derived Sequences. A targeted list of peptide sequences mapped specifically to the *NF1* gene or the synthetic minigene constructs. A Peptide FDR of < 0.1% was applied to ensure confident identification. All entries were filtered against the decoy and contaminants databases.

**Table S4. Comprehensive profiling of the HLA-A*02:01 immunopeptidome by IP-MS. Sheet 1:** A list of all identified HLA-A*02:01 binders. **Sheet 2:** Non-redundant HLA-A*02:01 binders and their length distribution. **Sheet 3:** A list of *NF1*-derived neoepitopes mapped to the *NF1* SAAVs. A Peptide FDR of < 0.1% was applied to ensure confident identification. All entries were filtered against the decoy and contaminants databases.

**Table S5. Direct mapping of HLA-A*02:01 binders to their cytosolic precursors.** A total of 265 9-mer HLA binders were identified with 100% sequence match to the endogenous peptide library. The corresponding precursor peptide sequences, length, UniProt IDs, and Gene IDs are provided in the third column.
