## Supplemental Figure 1 for "Large-Scale Assessment of *NF1* Single Amino Acid Variants as HLA Class I Neoantigens"

### NF1 Construct 1-8

#### Construct ID: NF1_C1

**Amino acid sequence**

**MRVTAPRTLILLLSGALALTETWAGSGGSGGGGSGGPGIDEETSEESLLTPTSPYLPALQSQLSITANGGSGGG
GSGGAIVSPYEAGILDKKPPPRIERSLKLMSKILQSIANHVLGGSGGGGSGGQRILYEYLAEASVVLPKVFPVV
HNLLDSKINTLLSLCQGGSGGGGSGGGKLAEHIEHEQQKLPAAILALEEDLKVFHNALKGGSGGGGSGGAGQPK
DTMRLDETMLAKQLLPEICHFLHTCREGGSLGGGGSGIVGIVAGLAVLAVVVIGAVVATVMCRRKSSGGKGGSY
SQAASSDSAQGSDVSLTA***

**Nucleotide sequence**

TGGCTAGTTAAGCTTGGATCCGCCACC**ATG**CGGGTGACTGCTCCACGGACACTGATTCTGCTTCTGTCAGGGGC
CTTGGCGCTCACCGAGACTTGGGCCGGATCAGGAGGCTCTGGAGGCGGAGGTTCTGGAGGCCCTGGAATTGATG
AAGAAACCAGTGAAGAATCCCTCCTGACTCCCACATCTCCTTACCTTCCTGCACTGCAGAGCCAGCTTAGTATC
ACTGCCAACGGAGGTAGCGGAGGCGGGGGTTCCGGCGGAGCCATTGTCTCACCGTATGAAGCAGGGATTTTAGA
TAAAAAGCCACCACCTAGAATCGAAAGGAGCTTGAAGTTAATGTCAAAGATACTTCAGAGTATTGCCAATCATG
TTCTCGGCGGGTCTGGAGGCGGGGGCTCAGGAGGCCAACGAATTCTTTATGAATACTTAGCAGAGGCCAGTGTT
GTGTTGCCCAAAGTCTTTCCTGTTGTGCATAATTTGTTGGACTCTAAGATCAACACCCTGTTATCATTGTGCCA
AGGTGGGAGTGGGGGTGGGGGAAGCGGCGGCGGGAAACTGGCTGAGCACATAGAGCATGAACAACAGAAACTAC
CTGCTGCCATCTTGGCTTTAGAAGAGGACCTGAAGGTATTCCACAATGCTCTCAAGGGGGGGTCCGGAGGCGGC
GGCTCCGGGGGCGCTGGGCAACCAAAGGACACAATGAGATTAGATGAAACGATGCTGGCCAAACAGTTGCTGCC
AGAAATCTGCCATTTTCTTCACACCTGTCGTGAAGGTGGATCTCTCGGAGGAGGAGGGAGCGGCATCGTCGGCA
TCGTCGCAGGCCTTGCGGTTCTGGCGGTCGTTGTCATCGGAGCAGTTGTGGCTACTGTCATGTGCAGGCGCAAA
AGCAGCGGCGGCAAAGGAGGTAGTTATTCACAGGCCGCCTCATCCGACAGTGCCCAAGGCTCCGATGTCTCACT
CACCGCG**TGA**ATTCACCATTGAGTTTAAA

#### Construct ID: NF1_C2

**Amino acid sequence**

**MRVTAPRTLILLLSGALALTETWAGSGGSGGGGSGGKVIGRMCKIIDKTYLSPTPTLEQHLMWDDIAILGGSGG
GGSGGLPGIDEETSEESLLTPTSPYPPALQRQLSITANLNLSNGGSGGGGSGGYLPGIDEETSEEALLTPTSPY
PPALQSQLSITANLNLSGGSGGGGSGGLPLQPEEGDGVELMEAESQLFLKYFTLFMNLLNDGGSGGGGSGGQRI
LYEYLAEASVVFPKVFPVVHDLLDSKINTLLSLCQGGSLGGGGSGIVGIVAGLAVLAVVVIGAVVATVMCRRKS
SGGKGGSYSQAASSDSAQGSDVSLTA***

**Nucleotide sequence**

TGGCTAGTTAAGCTTGGATCCGCCACC**ATG**CGGGTGACTGCTCCACGGACACTGATTCTGCTTCTGTCAGGGGC
CTTGGCGCTCACCGAGACTTGGGCCGGATCAGGAGGCTCTGGAGGCGGAGGTTCTGGAGGCAAGGTTATTGGAA
GGATGTGCAAAATAATTGACAAGACATACTTATCTCCAACTCCTACTTTAGAACAACATCTTATGTGGGATGAT
ATTGCTATTTTAGGAGGTAGCGGAGGCGGGGGTTCCGGCGGACTGCCTGGAATTGATGAAGAAACCAGTGAAGA
ATCCCTCCTGACTCCCACATCTCCTTACCCTCCTGCACTGCAGAGGCAGCTTAGTATCACTGCCAACCTTAACC
TTTCTAATGGCGGGTCTGGAGGCGGGGGCTCAGGAGGCTACCTGCCTGGAATTGATGAAGAAACCAGTGAAGAA
GCCCTCCTGACTCCCACATCTCCTTACCCTCCTGCACTGCAGAGCCAGCTTAGTATCACTGCCAACCTTAACCT
TTCTGGTGGGAGTGGGGGTGGGGGAAGCGGCGGCCTCCCTCTGCAGCCTGAAGAAGGAGATGGTGTGGAATTGA

TGGAAGCCGAATCACAGTTATTTCTTAAATACTTCACATTATTTATGAACCTTTTGAATGACGGGGGGTCCGGA
GGCGGCGGCTCCGGGGGCCAACGAATCCTTTATGAATACTTAGCAGAGGCCAGTGTTGTGTTTCCCAAAGTCTT
TCCTGTTGTGCATGATTTGTTGGACTCTAAGATCAACACCCTGTTATCATTGTGCCAAGGTGGATCTCTCGGAG
GAGGAGGGAGCGGCATCGTCGGCATCGTCGCAGGCCTTGCGGTTCTGGCGGTCGTTGTCATCGGAGCAGTTGTG
GCTACTGTCATGTGCAGGCGCAAAAGCAGCGGCGGCAAAGGAGGTAGTTATTCACAGGCCGCCTCATCCGACAG
TGCCCAAGGCTCCGATGTCTCACTCACCGCG**TGA**ATTCACCATTGAGTTTAAA

#### Construct ID: NF1_C3

**Amino acid sequence**

**MRVTAPRTLILLLSGALALTETWAGSGGSGGGGSGGYLSQLIILDTLENCLAGQPKDTMRLDETMLVKQLLPEI
GGSGGGGSGGCRQAQTKLEVALYMFPWNPDTEAVLVAMSCFRHLCEEAGGSGGGGSGGSATGGLGSIKAEVMAH
TAVALASGNVKLVSSKVIGGSGGGGSGGLRIFNDKSPEEVCMAIRNPLEWHCKQMDHFVGGGSGGGGSGGELSQ
PDSIPQHTNIRPKDVPGTLLNIALLNLGSSDPSLGGSLGGGGSGIVGIVAGLAVLAVVVIGAVVATVMCRRKSS
GGKGGSYSQAASSDSAQGSDVSLTA***

**Nucleotide sequence**

TGGCTAGTTAAGCTTGGATCCGCCACC**ATG**CGGGTGACTGCTCCACGGACACTGATTCTGCTTCTGTCAGGGGC
CTTGGCGCTCACCGAGACTTGGGCCGGATCAGGAGGCTCTGGAGGCGGAGGTTCTGGAGGCTATCTCTCTCAGT
TGATTATATTGGATACACTGGAAAATTGTCTTGCTGGGCAACCAAAGGACACAATGAGATTAGATGAAACGATG
CTGGTCAAACAGTTGCTGCCAGAAATCGGAGGTAGCGGAGGCGGGGGTTCCGGCGGATGCCGACAAGCCCAGAC
CAAACTAGAAGTGGCCCTGTACATGTTTCCGTGGAACCCTGACACTGAAGCTGTTCTGGTTGCCATGTCCTGTT
TCCGCCACCTCTGTGAGGAAGCAGGCGGGTCTGGAGGCGGGGGCTCAGGAGGCAGTGCAACAGGTGGCTTGGGA
TCAATAAAAGCTGAGGTGATGGCACATACTGCTGTAGCTTTGGCTTCTGGAAATGTGAAATTGGTTTCAAGCAA
GGTTATTGGTGGGAGTGGGGGTGGGGGAAGCGGCGGCCTCCGTATATTCAATGACAAGAGTCCAGAGGAAGTAT
GTATGGCAATCCGGAATCCTCTGGAGTGGCACTGCAAGCAAATGGATCATTTTGTTGGAGGGGGGTCCGGAGGC
GGCGGCTCCGGGGGCGAACTGTCACAGCCCGACTCTATCCCCCAACACACCAATATTCGGCCAAAAGATGTCCC
TGGGACACTGCTCAATATCGCATTACTTAATTTAGGCAGTTCTGACCCGAGTTTAGGTGGATCTCTCGGAGGAG
GAGGGAGCGGCATCGTCGGCATCGTCGCAGGCCTTGCGGTTCTGGCGGTCGTTGTCATCGGAGCAGTTGTGGCT
ACTGTCATGTGCAGGCGCAAAAGCAGCGGCGGCAAAGGAGGTAGTTATTCACAGGCCGCCTCATCCGACAGTGC
CCAAGGCTCCGATGTCTCACTCACCGCG**TGA**ATTCACCATTGAGTTTAAA

#### Construct ID: NF1_C4

**Amino acid sequence**

**MRVTAPRTLILLLSGALALTETWAGSGGSGGGGSGGLSQPDSIPQHTKIRPKNVPGTLLNIALLNLGSSDPSLR
GGSGGGGSGGPSLRSAAYNLLCVLTCTFNLKIEGQLLETSGLGGSGGGGSGGETRSYKYLLLSMVKPIHADPKL
LLCNPRKQGPETQGSTGGSGGGGSGGLPLQPEEGDGVELMEAISQLFLKYFTLFMNLLNDGGSGGGGSGGNTEA
WEDTHAKWEQATKPILNYPKAKMEDGQAGGSLGGGGSGIVGIVAGLAVLAVVVIGAVVATVMCRRKSSGGKGGS
YSQAASSDSAQGSDVSLTA***

**Nucleotide sequence**

TGGCTAGTTAAGCTTGGATCCGCCACC**ATG**CGGGTGACTGCTCCACGGACACTGATTCTGCTTCTGTCAGGGGC
CTTGGCGCTCACCGAGACTTGGGCCGGATCAGGAGGCTCTGGAGGCGGAGGTTCTGGAGGCCTGTCACAGCCCG
ACTCTATCCCCCAACACACCAAGATTCGGCCAAAAAATGTCCCTGGGACACTGCTCAATATCGCATTACTTAAT
TTAGGCAGTTCTGACCCGAGTTTACGGGGAGGTAGCGGAGGCGGGGGTTCCGGCGGACCGAGTTTACGGTCAGC
TGCCTATAATCTTCTGTGTGTCTTAACTTGTACCTTTAATTTAAAAATCGAGGGCCAGTTACTAGAGACATCAG
GTTTAGGCGGGTCTGGAGGCGGGGGCTCAGGAGGCGAGACAAGAAGCTATAAGTATCTTCTCTTGTCCATGGTG
AAACCAATTCATGCAGATCCAAAGCTCTTGCTTTGTAATCCAAGAAAACAGGGGCCCGAAACCCAAGGCAGTAC
AGGTGGGAGTGGGGGTGGGGGAAGCGGCGGCCTCCCTCTGCAGCCTGAAGAAGGAGATGGTGTGGAATTGATGG
AAGCCATATCACAGTTATTTCTTAAATACTTCACATTATTTATGAACCTTTTGAATGACGGGGGGTCCGGAGGC
GGCGGCTCCGGGGGCAACACTGAGGCTTGGGAAGATACACATGCAAAATGGGAACAAGCAACAAAGCCAATCCT
TAACTATCCAAAAGCCAAAATGGAAGATGGCCAGGCTGGTGGATCTCTCGGAGGAGGAGGGAGCGGCATCGTCG
GCATCGTCGCAGGCCTTGCGGTTCTGGCGGTCGTTGTCATCGGAGCAGTTGTGGCTACTGTCATGTGCAGGCGC
AAAAGCAGCGGCGGCAAAGGAGGTAGTTATTCACAGGCCGCCTCATCCGACAGTGCCCAAGGCTCCGATGTCTC
ACTCACCGCG**TGA**ATTCACCATTGAGTTTAAA

#### Construct ID: NF1_C5

##### Amino acid sequence

**MRVTAPRTLILLLSGALALTETWAGSGGSGGGGSGGGFSKSSIELKHLCLQYMTPWLSNLVRFCKHNDDGGSGG
GGSGGMLSFNNSLDVAAHLPYLFHVVTFLLATGPLSLRASTHGGGSGGGGSGGLPGIDEETSEESFLTPTSPYP
PALQSQLSITANLNLSNGGSGGGGSGGAGLPLQPEEGDGVELMEAKSQLFRKYFTLFMNLLNDCSGGSGGGGSG
GRISPHNNQHFKIYLAQNSPSTFHYVLVNSLHRIITNSAGGSLGGGGSGIVGIVAGLAVLAVVVIGAVVATVMC
RRKSSGGKGGSYSQAASSDSAQGSDVSLTA***

##### Nucleotide sequence

TGGCTAGTTAAGCTTGGATCCGCCACC**ATG**CGGGTGACTGCTCCACGGACACTGATTCTGCTTCTGTCAGGGGC
CTTGGCGCTCACCGAGACTTGGGCCGGATCAGGAGGCTCTGGAGGCGGAGGTTCTGGAGGCGGATTTAGCAAAT
CTAGTATTGAATTGAAACACCTTTGTTTGCAATACATGACTCCATGGCTGTCAAATCTAGTTCGTTTTTGCAAG
CATAATGATGATGGAGGTAGCGGAGGCGGGGGTTCCGGCGGAATGCTGTCCTTCAACAATTCCCTTGATGTGGC
AGCTCATCTTCCCTACCTCTTCCACGTTGTTACTTTCTTATTAGCCACAGGTCCGCTCTCCCTTAGAGCTTCCA
CACATGGAGGCGGGTCTGGAGGCGGGGGCTCAGGAGGCCTGCCTGGAATTGATGAAGAAACCAGTGAAGAATCC
TTCCTGACTCCCACATCTCCTTACCCTCCTGCACTGCAGAGCCAGCTTAGTATCACTGCCAACCTTAACCTTTC
TAATGGTGGGAGTGGGGGTGGGGGAAGCGGCGGCGCTGGTCTCCCTCTGCAGCCTGAAGAAGGAGATGGTGTGG
AATTGATGGAAGCCAAATCACAGTTATTTCGTAAATACTTCACATTATTTATGAACCTTTTGAATGACTGCAGT
GGGGGGTCCGGAGGCGGCGGCTCCGGGGGCCGTATAAGCCCTCACAACAACCAACACTTTAAGATCTACCTGGC
TCAGAAcTCACCTTCTACATTTCACTATGTGCTGGTAAATTCACTCCATCGAATCATCACCAATTCCGCAGGTG
GATCTCTCGGAGGAGGAGGGAGCGGCATCGTCGGCATCGTCGCAGGCCTTGCGGTTCTGGCGGTCGTTGTCATC
GGAGCAGTTGTGGCTACTGTCATGTGCAGGCGCAAAAGCAGCGGCGGCAAAGGAGGTAGTTATTCACAGGCCGC
CTCATCCGACAGTGCCCAAGGCTCCGATGTCTCACTCACCGCG**TGA**ATTCACCATTGAGTTTAAA

#### Construct ID: NF1_C6

##### Amino acid sequence

**MRVTAPRTLILLLSGALALTETWAGSGGSGGGGSGGRYMLMLSFNNSLDVAAHPPYLFHVVTFLVATGPGGSGG
GGSGGILICRNKFLLKNKQADRSSCHFLLFCGVGCDIPSSGNTGGSGGGGSGGLSSTEILKWLREILICRNNFL
LKNKQADRSSCHFGGSGGGGSGGSVSESNVLLDEEVLTDPKIQVLLLTVLATLVKYTTDEFGGSGGGGSGGDVP
GTLLNIALLNLDSSDPSLRSAAYNLLCALTGGSLGGGGSGIVGIVAGLAVLAVVVIGAVVATVMCRRKSSGGKG
GSYSQAASSDSAQGSDVSLTA***

#### Nucleotide sequence

TGGCTAGTTAAGCTTGGATCCGCCACC**ATG**CGGGTGACTGCTCCACGGACACTGATTCTGCTTCTGTCAGGGGC
CTTGGCGCTCACCGAGACTTGGGCCGGATCAGGAGGCTCTGGAGGCGGAGGTTCTGGAGGCCGCTACATGCTGA
TGCTGTCCTTCAACAATTCCCTTGATGTGGCAGCTCATCCTCCCTACCTCTTCCACGTTGTTACTTTCTTAGTA
GCCACAGGTCCGGGAGGTAGCGGAGGCGGGGGTTCCGGCGGAATATTGATCTGCAGGAATAAATTTCTTCTTAA
AAATAAGCAGGCAGATAGAAGTTCCTGTCACTTTCTCCTTTTTTGCGGGGTAGGATGTGATATTCCTTCTAGTG
GAAATACCGGCGGGTCTGGAGGCGGGGGCTCAGGAGGCCTTAGTAGCACAGAAATTCTCAAGTGGTTGCGGGAA
ATATTGATCTGCAGGAATAACTTTCTTCTTAAAAATAAGCAGGCAGATAGAAGTTCCTGTCACTTTGGTGGGAG
TGGGGGTGGGGGAAGCGGCGGCTCAGTGTCTGAATCAAATGTTCTCTTGGATGAAGAAGTACTTACTGATCCGA
AGATCCAGGTGCTGCTTCTTACTGTTCTAGCTACACTGGTAAAATATACCACAGATGAGTTTGGGGGGTCCGGA
GGCGGCGGCTCCGGGGGCGATGTCCCTGGGACACTGCTCAATATCGCATTACTTAATTTAGACAGTTCTGACCC
GAGTTTACGGTCAGCTGCCTATAATCTTCTGTGTGCCTTAACTGGTGGATCTCTCGGAGGAGGAGGGAGCGGCA
TCGTCGGCATCGTCGCAGGCCTTGCGGTTCTGGCGGTCGTTGTCATCGGAGCAGTTGTGGCTACTGTCATGTGC
AGGCGCAAAAGCAGCGGCGGCAAAGGAGGTAGTTATTCACAGGCCGCCTCATCCGACAGTGCCCAAGGCTCCGA
TGTCTCACTCACCGCG**TGA**ATTCACCATTGAGTTTAAA

#### Construct ID: NF1_C7

#### Amino acid sequence

**MRVTAPRTLILLLSGALALTETWAGSGGSGGGGSGGTSLETVTEALLEIMEACMREIPTCKWLDQWTELGGSGG
GGSGGLMHSIGLGYHKDLQTRATFMKVLTKILQQGTEFDTLAEGGSGGGGSGGPTSDAVNHSLSFISDGNVLVL
HRLLWNNQEKIGGGSGGGGSGGHAIQIKTKLCQLIEVMMARRDDLSFCQEMKFRNKMVEYGGSGGGGSGGLSST
EILKWLREILICRNKFLLKNNQADRSSCHFLLFGGSLGGGGSGIVGIVAGLAVLAVVVIGAVVATVMCRRKSSG
GKGGSYSQAASSDSAQGSDVSLTA***

#### Nucleotide sequence

TGGCTAGTTAAGCTTGGATCCGCCACC**ATG**CGGGTGACTGCTCCACGGACACTGATTCTGCTTCTGTCAGGGGC
CTTGGCGCTCACCGAGACTTGGGCCGGATCAGGAGGCTCTGGAGGCGGAGGTTCTGGAGGCACATCCTTGGAAA
CAGTCACAGAAGCTTTGTTGGAGATCATGGAGGCATGCATGAGAGAGATTCCAACGTGCAAGTGGCTGGACCAG
TGGACAGAACTAGGAGGTAGCGGAGGCGGGGGTTCCGGCGGACTCATGCACTCCATAGGCTTAGGTTACCACAA
GGATCTCCAGACAAGAGCTACATTTATGAAAGTTCTGACAAAAATCCTTCAACAAGGCACAGAATTTGACACAC
TTGCAGAAGGCGGGTCTGGAGGCGGGGGCTCAGGAGGCCCTACAAGTGATGCAGTAAATCATAGTCTTTCCTTC
ATAAGTGACGGCAATGTGCTTGTTTTACATCGTCTACTCTGGAACAATCAGGAGAAAATTGGGGGTGGGAGTGG
GGGTGGGGGAAGCGGCGGCCATGCAATTCAAATAAAAACGAAACTGTGTCAATTAATTGAAGTAATGATGGCAA
GGAGAGATGACCTCTCATTTTGCCAAGAGATGAAATTTAGGAATAAGATGGTAGAATACGGGGGGTCCGGAGGC
GGCGGCTCCGGGGGCCTTAGTAGCACAGAAATTCTCAAGTGGTTGCGGGAAATATTGATCTGCAGGAATAAATT
TCTTCTTAAAAATAACCAGGCAGATAGAAGTTCCTGTCACTTTCTCCTTTTTGGTGGATCTCTCGGAGGAGGAG
GGAGCGGCATCGTCGGCATCGTCGCAGGCCTTGCGGTTCTGGCGGTCGTTGTCATCGGAGCAGTTGTGGCTACT

GTCATGTGCAGGCGCAAAAGCAGCGGCGGCAAAGGAGGTAGTTATTCACAGGCCGCCTCATCCGACAGTGCCCA
AGGCTCCGATGTCTCACTCACCGCG**TGA**ATTCACCATTGAGTTTAAA

**Construct ID: NF1_C8**

**Amino acid sequence**

**MRVTAPRTLILLLSGALALTETWAGSGGSGGGGSGGLQYINVDCAKLKRLLKKTAFKFKALKKVAQLAVINSLG
GSGGGGSGGSLLAGLPLQPEEEDGVELMEAKSQLFLKYFTLFMNLLNGGSGGGGSGGHQECEAIVQSIIHMRTR
WELSQPDSIPQHTKIGGSGGGGSGGILICRNKFLLKNKQADRSSFHFLLFYGVGCDIPSSGGGSGGGGSGGEGY
LAATYPTVGQISPRARKSMSLDMGQPSQANTKKLGGSLGGGGSGIVGIVAGLAVLAVVVIGAVVATVMCRRKSS
GGKGGSYSQAASSDSAQGSDVSLTA***

**Nucleotide sequence**

TGGCTAGTTAAGCTTGGATCCGCCACC**ATG**CGGGTGACTGCTCCACGGACACTGATTCTGCTTCTGTCAGGGGC
CTTGGCGCTCACCGAGACTTGGGCCGGATCAGGAGGCTCTGGAGGCGGAGGTTCTGGAGGCTTACAGTATATCA
ATGTGGATTGTGCAAAATTAAAACGACTCCTGAAGAAAACAGCATTTAAATTTAAAGCCCTAAAGAAGGTTGCG
CAGTTAGCAGTTATAAATAGCCTGGGAGGTAGCGGAGGCGGGGGTTCCGGCGGATCACTTCTAGCTGGTCTCCC
TCTGCAGCCTGAAGAAGAAGATGGTGTGGAATTGATGGAAGCCAAATCACAGTTATTTCTTAAATACTTCACAT
TATTTATGAACCTTTTGAATGGCGGGTCTGGAGGCGGGGGCTCAGGAGGCCACCAGGAGTGTGAAGCCATTGTC
CAGTCTATCATTCATATGCGGACCCGCTGGGAACTGTCACAGCCCGACTCTATCCCCCAACACACCAAGATTGG
TGGGAGTGGGGGTGGGGGAAGCGGCGGCATATTGATCTGCAGGAATAAATTTCTTCTTAAAAATAAGCAGGCAG
ATAGAAGTTCCTTTCACTTTCTCCTTTTTTACGGGGTAGGATGTGATATTCCTTCTAGTGGAGGGGGGTCCGGA
GGCGGCGGCTCCGGGGGCGAAGGATACCTTGCAGCCACCTATCCAACTGTCGGCCAGATCAGTCCCCGAGCCAG
GAAATCCATGAGCCTGGACATGGGGCAACCTTCTCAGGCCAACACTAAGAAGTTGGGTGGATCTCTCGGAGGAG
GAGGGAGCGGCATCGTCGGCATCGTCGCAGGCCTTGCGGTTCTGGCGGTCGTTGTCATCGGAGCAGTTGTGGCT
ACTGTCATGTGCAGGCGCAAAAGCAGCGGCGGCAAAGGAGGTAGTTATTCACAGGCCGCCTCATCCGACAGTGC
CCAAGGCTCCGATGTCTCACTCACCGCG**TGA**ATTCACCATTGAGTTTAAA

**Color Key:**

**5′ linker:** TGGCTAGTTAAGCTTGGATCC
**3′ linker:** ATTCACCATTGAGTTTAAA

**GCCACC: Kozak sequence** (GCCACC**ATG**G)

**MRVTAPRTLILLLSGALALTETWAGS: MHC class I signal peptide (secretion signal)**

**GGSGGGGSGG: GS linker**

**GGSLGGGGSG: GS linker**

**IVGIVAGLAVLAVVVIGAVVATVMCRRKSSGGKGGSYSQAASSDSAQGSDVSLTA*: MHC class I trafficking signal (MITD)**
